# Fluorescently Labeled Gradient Hydrogels Reveal Matrix-Dependent Cell Responses to Substrate Stiffness

**DOI:** 10.1101/2025.09.23.677581

**Authors:** Shin Wei Chong, Li Liu, Daryan Kempe, Yingqi Zhang, Kourosh Kalantar-Zadeh, Marcela M.M. Bilek, Lining Arnold Ju, Maté Biro, Daniele Vigolo

## Abstract

Microfabricated stiffness gradient hydrogels hold significant value for advancing mechanobiology, tissue engineering, and *in vitro* tissue models. However, it remains challenging to design these materials given their broad processing parameter space. The continuum of stiffness values also makes it difficult to precisely correlate the local substrate properties and observed biological responses, often relying on cumbersome characterization methods such as atomic force microscopy. To address these bottlenecks, we present a straightforward thermophoresis-based fabrication strategy to pattern stiffness gradients in fluorescein isothiocyanate-labeled hydrogel network, which displays a polymer concentration-dependent fluorescence readout. This approach enables quantitative assessment of the gradient formation process and contactless stiffness mapping *via* standard microscopy imaging. Using gelatin methacryloyl and Gellan gum as model systems, it is shown that substrate stiffness and extracellular matrix protein composition work together to affect 3T3-L1 fibroblast cell morphology and migration, with the underlying hydrogel type also affecting the outcome. By offering a simple and reliable approach for characterizing stiffness gradient hydrogels, this work advances the thermophoretic fabrication platform, opening avenues for new biomaterial systems for understanding and controlling the cell-material interplay.

## 1. Introduction

Microscale stiffness gradients are increasingly better understood as an essential feature with important biological functions in living tissues.^[1,2]^ This property of the extracellular matrix (ECM) has provided motivation for the development of materials that can mimic the pathophysiological mechanical landscape, leading to the creation of microfabricated stiffness gradient hydrogels for a variety of applications, such as cell-substrate interaction studies in mechanobiology,^[3,4]^ *in vitro* models of development and disease,^[5–7]^ and tissue engineering.^[8,9]^ Despite substantial progress in fabrication techniques, achieving effective design, validation, and optimization of microscale stiffness gradients continues to be a major challenge. The generation of a continuous stiffness gradient requires precise control of the polymer concentration (e.g., two-step polymerization,^[10]^ microfluidic mixing,^[11,12]^ thermophoresis^[13]^) or crosslinking density (e.g., graded exposure,^[14,15]^ freeze-thaw^[16]^). Once fabricated, individual gradient hydrogels should be comprehensively characterized to ensure the reliability and reproducibility for cell experiments, typically requiring specialized tools such as atomic force microscopy (AFM).^[17]^ Key challenges in these workflows include efficiently identifying optimal conditions to achieve the desired gradient features, developing scalable method for high-throughput experimentation, and establishing direct spatial correlation of local hydrogel stiffness with cellular responses. It is therefore important to address these practical bottlenecks, as they directly affect both the predictive value of *in vitro* stiffness gradient hydrogel studies and their broader technological adoption.

Existing workflows largely rely on experimental optimization and testing various parameter combinations for uncovering material performance insights.^[18]^ However, regardless of the fabrication method, there are multiple variables that can affect the stiffness values along the gradient gel. For example, in the context of a thermophoresis-based process (i.e., temperature-field-activated polymer network migration strategy),^[19,20]^ these include the polymer physicochemical properties, base polymer and crosslinker concentration, applied temperature field strength, gel size, and the temperature field induction time.^[21–24]^ To date, most gradient hydrogel platforms have been advanced through iterative processes of trial-and-error, but there are limitations. Humans are generally not equipped to synthesize high-dimensionality data, making processes with multiple inputs/outputs challenging to analyze.^[25]^ In this regard, the time and effort associated with manual one-hydrogel-at-a-time fabrication and characterization can be burdensome. Moreover, the lack of a predictive design framework means that the fabrication optimization must be repeated for each new gradient configuration, limiting experimental flexibility.^[20]^

Among the suite of techniques for measuring the stiffness (i.e., Young’s modulus) of soft hydrogels, AFM is the most established, offering sensitivity at the scale relevant to how single cells ‘feel’ their surrounding matrix, and thus remains the workhorse for mechanobiology-related studies.^[26,27]^ However, AFM is inherently low throughput (≈45 min per gradient sample), labor-intensive, and often not readily accessible to standard laboratories. To overcome such challenges, fluorescence-based strategies have emerged as promising alternatives for stiffness gradient characterization.^[28]^ For example, a recent study has demonstrated that the diffusion of fluorescein within a polyacrylamide precursor mix could be directly correlated with the resulting stiffness gradient, enabling fluorescence intensity to be used as an optical readout of local hydrogel stiffness.^[29]^ While straightforward, this method can be highly dependent on the imaging setup and is susceptible to photobleaching. In another approach, fluorescent microbeads have been incorporated during hydrogel fabrication as a more stable, time-invariant proxy of substrate stiffness based on the local bead density.^[4,30]^ However, this approach did not provide a direct view of the polymer network. Based on such shortcomings, there is a strong case for developing alternative strategies based on fluorescence imaging to characterize stiffness gradient hydrogels, where the polymers are directly labeled and tracked,^[31]^ thereby enabling precise correlation between the polymer composition, hydrogel mechanics, and cell behavior for comprehensive investigations of cell-material interplay.

This study presents a strategy leveraging well-established fluorescein isothiocyanate (FITC)-labeling chemistry to allow real-time monitoring of the stiffness gradient formation and material-intrinsic indication of the local hydrogel stiffness that is readily integrated in existing experiment workflows. Focusing on the temperature-field-driven thermophoretic fabrication method, it is demonstrated that preliminary experimental exploration of the process dynamics enabled the identification of empirical relationships between the material system, process design, and resulting gradient outcomes. Such exploration streamlines the material optimization by eliminating arduous cycles of gradient gel fabrication and assessment, and also carries significant potential toward predictive design frameworks for materials development. Using this approach, the resulting stiffness gradient hydrogels exhibited stable fluorescence signal enabling high-resolution and precise optical readout of local hydrogel stiffness. Furthermore, its versatility and potential value of the developed method for mechanobiology are shown by investigating a series of gelatin methacryloyl (GelMA) and Gellan gum gradient gels, evaluating the complex interplay between stiffness, ECM protein coating, and underlying substrate type on 3T3-L1 fibroblast mechanosensing, relevant to the design of biomaterials for programmable cell behavior.

## 2. Results

### 2.1. Preparation of Fluorescently Labeled Hydrogels *via* FITC Conjugation

The development of fluorescently labeled hydrogels was based on covalent conjugation of FITC fluorophores directly onto the polymer backbone. FITC was chosen due to its low cost, accessibility, and the established reactivity of its isothiocyanate group (–N=C=S) to form stable bonds with primary amines (–NH_2_) or hydroxyl (–OH) groups (**Figure 1A**).^[31,32]^ These functional groups are commonly found across both natural and synthetic polymers, which illustrates the versatility of the fluorescence strategy. We focus our study on carbohydrate-based Gellan gum and chemically modified collagen-based GelMA, both of which are widely used ECM mimics in the broad fields of biofabrication and tissue engineering.^[33–35]^ To ensure that the fluorescence signal originates specifically from the labeled polymers, unbound FITC molecules were removed through dialysis after the labeling procedure. In both cases of FITC-GelMA (F-GM) and FITC-Gellan gum (F-GG), the final product was a yellow fibrous foam (**Figure 1B, 1C**). Importantly, low degrees of FITC substitution were employed to provide robust fluorescence signals without affecting the gelation behavior (**Figure 1D, 1E**). The substituted FITC content as mass fraction was ::0.01 for F-GM and ::0.004 for F-GG, evaluated by fluorescence intensity measurements using a microplate reader, and which fall within the typical ranges reported in other studies (**Supporting Information, Figure S1**).^[36,37]^

**Figure 1:**
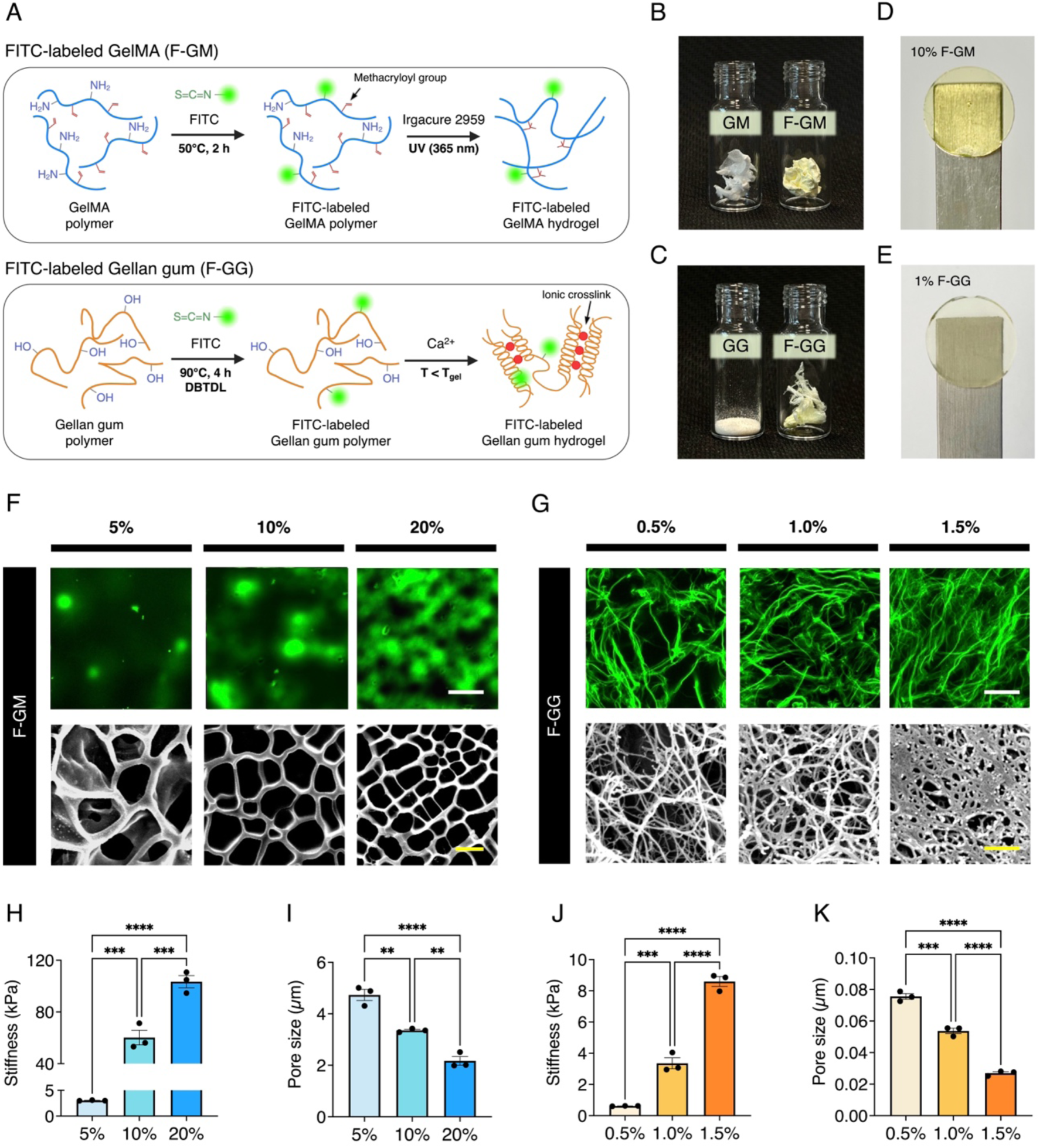
Preparation, structure, and mechanical characterization of FITC-labeled GelMA (F-GM) and Gellan gum (F-GG). (A) Schematic of the synthesis and formation of F-GM (top) and F-GG (bottom) hydrogels. (B) Photograph of lyophilized GelMA before and after fluorescent labeling. (C) Photograph of Gellan gum powder as received commercially and the lyophilized form after fluorescent labeling. (D) Hydrogels were formed by standard photopolymerization for F-GM and (E) ionic crosslinking for F-GG. (F) Representative confocal (top) and SEM (bottom) images of F-GM hydrogels prepared at varied concentrations (5 wt%, 10 wt%, 20 wt%). Scale bar: 25 µm (confocal), 5 µm (SEM). The corresponding hydrogel Young’s modulus and pore size are shown in (H) and (I), respectively (n = 3 independent samples; mean ± SEM). (G) Representative confocal (top) and SEM (bottom) images of F-GG hydrogels prepared at varied concentrations (0.5 wt%, 1.0 wt%, 1.5 wt%). Scale bar: 25 µm (confocal), 0.5 µm (SEM). The corresponding hydrogel stiffness and pore size are shown in (J) and (K), respectively (n = 3 independent samples; mean ± SEM). Quantification of pore sizes were performed on SEM images. * (p < 0.05), ** (p < 0.01), *** (p < 0.005), ns (not significant).

In preliminary characterization studies, uniform stiffness gels were fabricated with 5 wt% to 20 wt% F-GM (**Figure 1F**) or 0.5 wt% to 1.5 wt% F-GG (**Figure 1G**). Through AFM nanoindentation, it was found that the Young’s modulus of the resulting gels increased proportionally to the polymer concentration over these ranges (**Figure 1H, 1J**). Moreover, structural visualization of the hydrogel network was performed using confocal fluorescence microscopy and scanning electron microscopy (SEM) (**Figure 1F, 1G**). The results showed, as expected, a porous mesh morphology in F-GM and a fibrous architecture in F-GG, confirming that the addition of FITC does not disrupt proper hydrogel network formation. The confocal images exhibited a clear decrease in dark regions with increasing polymer concentration, suggesting that the dark regions were voids within the hydrogel network. A higher amount of polymer content leads to denser crosslinking and, thus, smaller pore sizes, consistent with the SEM analysis (**Figure 1I, 1K**). Note that while fluorescent labeling did not affect the mechanical properties of F-GM hydrogels, the F-GG hydrogels were generally softer than the unmodified Gellan gum (**Supplementary Information, Figure S2**). A similar behavior has also been observed in a previous report, suggesting that the molecular size of FITC could potentially alter the Gellan chain interactions.^[38]^ Nevertheless, both F-GM and F-GG supported repeatable hydrogel formation that retained structural integrity and stable fluorescence signal under cell culture conditions, demonstrating their feasibility for tunable stiffness gradient formation. Furthermore, the range of stiffness developed across both F-GM (≈3 to 100 kPa) and F-GG (≈0.5 to 9 kPa) was broadly relevant to the mechanical environments encountered by fibroblasts in healthy and fibrotic tissues.^[39]^ Taken together, these results establish fluorescent GelMA and Gellan gum with tunable stiffness and stable fluorescence, enabling the capacity to mechanically pattern gradient hydrogels for direct comparison of substrate type-dependent cellular stiffness responses.

### 2.2. Thermophoretic Polymers Migration and Experimental Optimization of Gradient Hydrogel Fabrication

The generation of gradient polymer concentration driven by an external temperature field is the key to thermophoretic fabrication of stiffness gradient hydrogels.^[19,20]^ To understand the material-specific thermophoretic behavior, a bespoke microfluidic device was designed for on-chip temperature control and direct observation of the polymer redistribution *via* standard fluorescence microscopy (**Figure 2A, 2B**).^[40]^ In this study, F-GM and F-GG exhibited a fluorescence intensity that positively correlated with the polymer concentration (**Figure 2C**). Thus, thermophoretic migration of the polymers in solution produced a spatial fluorescence pattern that varied over time, which can be analyzed to extract useful process information. It is worth noting that the absolute stiffness range and gradient strength, which are two crucial considerations when designing gradient hydrogels for cell culture experiments, are influenced by several factors, including the starting polymer concentration and applied temperature field strength and duration. Finding the optimal combination of these factors is therefore important to achieve consistent and reproducible stiffness gradients, as well as the desired biological outcomes.

**Figure 2:**
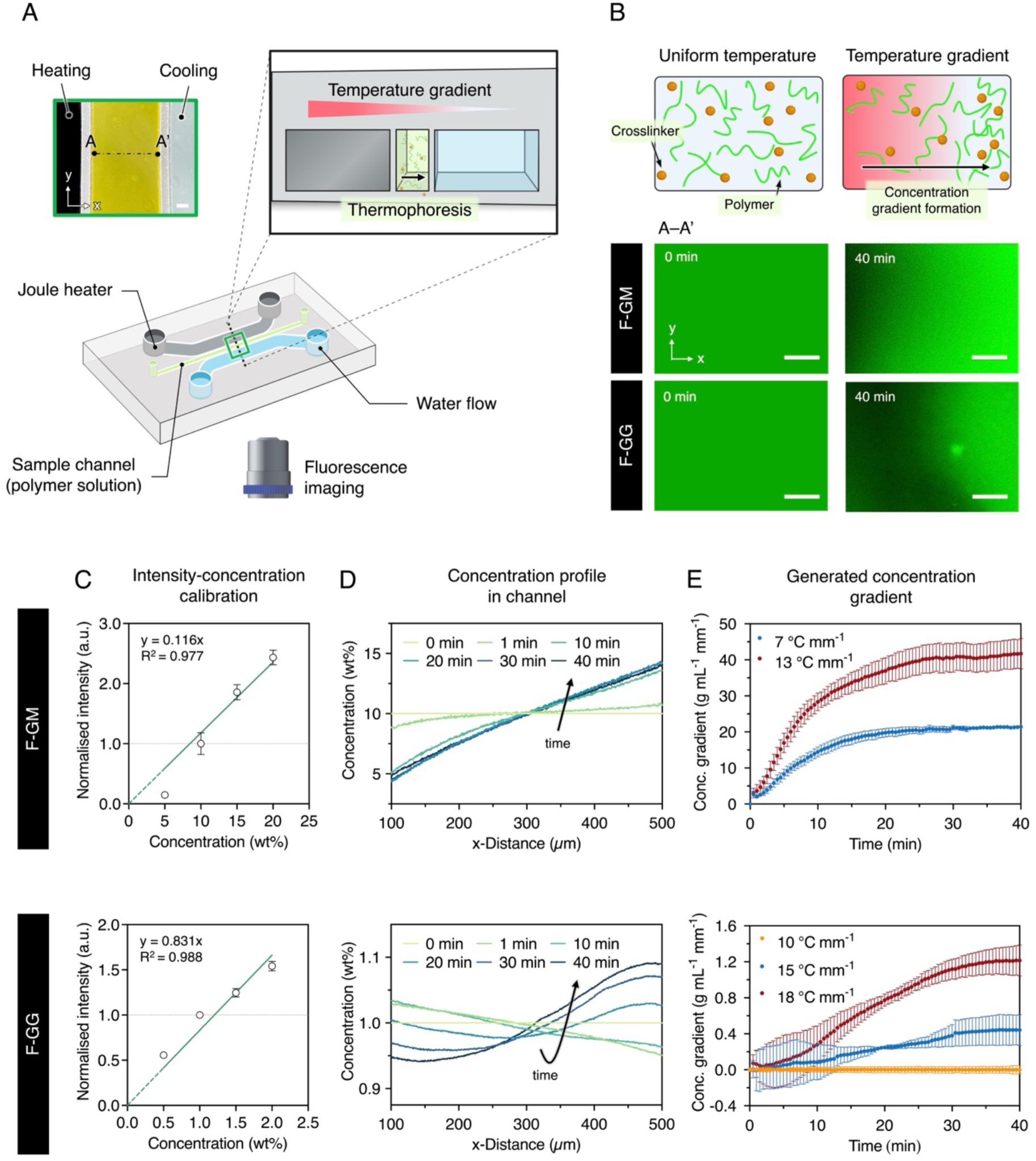
Fluorescence analysis of polymer gradient formation by thermophoresis. (A) Schematic of the experimental setup. The green bounding box shows a zoom-in optical image from the top view of the microfluidic device. The black bounding box illustrates the cross-section of the device. (B) Schematic of the mechanism of gradient formation in the presence of a temperature gradient, with representative fluorescent images demonstrating the accumulation of F-GM and F-GG polymer chains from hot to cold regions. Applied temperature gradient = 7 °C mm^-1^ (F-GM), 15 °C mm^-1^ (F-GG). (C) Correlation curve connecting F-GM (top) and F-GG (bottom) polymer concentration to fluorescence intensity (n = 3 independent samples; mean ± SEM). (D) Change in concentration profile for F-GM (top) and F-GG (bottom) over 40 min for the examples in (B). (E) The resulting concentration gradient as a function of time for F-GM (top) and F-GG (bottom) show a dependence on the applied temperature conditions (n = 3 independent experiments; mean ± SEM). Scale bars: 100 µm.

When working with a new hydrogel system, the concentration of the base solution is first selected to match the target stiffness range. The goal of the thermophoresis study is then to 1) establish the directionality of polymer migration, 2) assess the threshold temperature and duration to generate a discernable gradient distribution, and 3) examine how changes in the magnitude of applied temperature field affect the resulting polymer concentration gradient.

To demonstrate the workflow, this section is focused on experiments using 10 wt% F-GM and 1 wt% F-GG. As shown in **Figures 2B and 2D**, polymers within an initially homogenous system displayed directional drift within the first minute and progressively accumulated toward the cold region under constant temperature field application. The accumulation followed an exponential decay behavior characteristic of thermophoresis and plateaued at a concentration gradient that scales proportionally with the applied temperature gradient strength (**Figure 2E**).^[41]^ From these concentration gradient curves, it was also possible to determine the thermophoresis characteristic time, τ, which describes how quickly the system reaches equilibrium (**Supporting Information, Figure S3**). Generally, a stable F-GM concentration gradient was seen to be generated after applying a temperature field as low as 7 °C mm^-1^ for 30 min. On the other hand, F-GG showed a higher threshold temperature gradient (::15 °C mm^-1^) to induce a discernible thermophoresis effect. Furthermore, the time to reach steady state in the F-GG system was longer even at higher applied temperature conditions, which can be explained by the significantly larger molecular size of Gellan gum (::1000 kDa) compared to GelMA (::100 kDa).^[42,43]^ Importantly, these results provided mechanistic confirmation that the temperature-field-dependent manipulation could enable the universal fabrication of microscale gradient hydrogels by spatially encoding polymer distributions prior to crosslinking.

The process insights gained from the thermophoresis experiments are directly transferrable across different microconfinement setups and hydrogel dimensions. At a given thermal condition, the required process duration for gradient generation scales proportionally with the distance squared (i.e., 𝝉 ∝ 𝒅^𝟐^) corresponding to a diffusive behavior.^[19]^ In this study, findings from the initial thermophoresis characterization performed using a 600 µm-wide microchannel were successfully applied to fabricate 1000 µm-wide stiffness gradient hydrogels using an open microchamber system (**Figure 3A**). By eliminating the need for iterative trial-and-error of gradient hydrogel fabrication and stiffness measurements typical in traditional workflows, this process development strategy accelerates optimization, is designed to be accessible using standard laboratory equipment, and opens new physics-informed avenues for predictable gradient material design.

**Figure 3:**
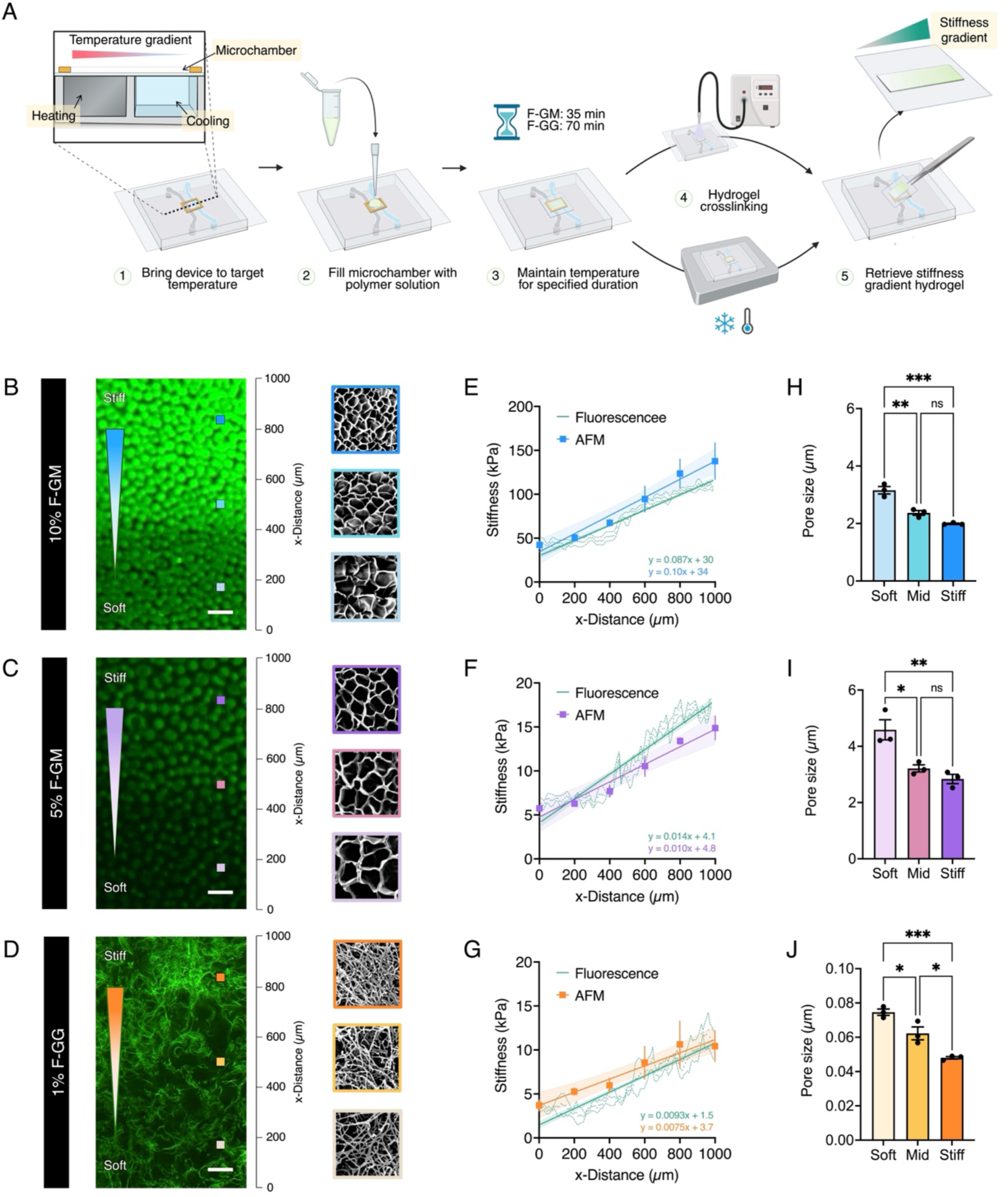
Fabrication and characterization of linear stiffness gradient hydrogels. (A) Graphical illustration of the stepwise fabrication procedure of gradient F-GM and F-GG hydrogels. The precursor solution is contained in an external microchamber and subjected to a defined heating pattern to create a continuous stiffness gradient upon hydrogel crosslinking. Representative fluorescent images of gradient gels fabricated using (B) 10 wt% F-GM, (C) 5 wt% F-GM, and (D) 1 wt% F-GG polymer systems. Scale bars: 100 µm. Colored squares indicate the approximate locations where SEM images were acquired, corresponding to stiff (top), moderately stiff (middle), and soft (bottom) regions of the gradient gels. Each square is 25 µm × 25 µm for F-GM samples and 2 µm × 2 µm for F-GG samples. The change in Young’s modulus of the gradient gels was measured by AFM nanoindentation and fluorescence mapping, yielding (E) steep gradient F-GM, (F) shallow gradient F-GM, and (G) shallow gradient F-GG hydrogels (n = 3 independent samples; mean ± SEM). Solid lines are simple linear regression plots with 95% confidence intervals shaded accordingly. Corresponding quantification of the pore size shows an inverse correlation with stiffness for (H) steep gradient F-GM, (I) shallow gradient F-GM, and (J) shallow gradient F-GG hydrogels (n = 3 independent samples; mean ± SEM). * (p < 0.05), ** (p < 0.01), *** (p < 0.005), ns (not significant).

### 2.3. Contactless Stiffness Mapping *via* Fluorescence

For studying cell-substrate interactions, the ability to directly correlate the fluorescence intensity with the mechanical properties of these FITC-labeled hydrogels endows another major advantage by enabling straightforward stiffness readouts *via* fluorescence visualization. To demonstrate their practicability, linear stiffness gradient hydrogels were fabricated using a second microfluidic system that we have developed recently.^[20]^ Using this new platform, the temperature gradient was imposed adjacently to an external sample microchamber, greatly simplifying setup and operation (**Figure 3A**, Steps 1-3). After that, the established gradient polymer network was permanently fixed by rapid crosslinking, resulting in a uniform stiffness gradient with the softest surface near the hottest region, to the stiffest near the coldest region (**Figure 3A**, Steps 4-5). Stiffness gradients with different slope profiles were produced in F-GM hydrogels using 10 wt% (stiffer, wide-range gradient; **Figure 3B**) or 5 wt% (softer, narrow-range gradient; **Figure 3C**) initial polymer concentration, mimicking pathological and physiological tissue conditions, respectively (**Supporting Information, Figure S4**).^[44]^ Furthermore, gradient F-GG hydrogels were produced using 1 wt% initial polymer concentration, which was intended to match the softer range gradient of the F-GM samples (**Figure 3D**).

Confocal scanning of the hydrogel samples consistently revealed a spatial gradient in fluorescence signal, indicating successful formation of a gradient gel network structure. Additionally, SEM images of the freeze-dried gradient hydrogels clearly exhibited a decrease in pore size from the darkest to the brightest regions, which confirmed that the gradient network was contributed by local variations in polymer distribution across the hydrogel (**Figure 3H, 3I, 3J**). The higher the fluorescence intensity, the higher the polymer concentration and, thus, mechanical stiffness. These variables can be connected using two correlation curves: fluorescence intensity as a function of polymer concentration (**Figure 2C**), and concentration versus stiffness as determined by AFM (**Figure 1H and 1J**). As such, by evaluating the relative intensity change with reference to the baseline (at known initial polymer concentration) and comparing against the established correlation curves, it is possible to infer the AFM-defined stiffness at any given point within the hydrogel (see **Supplementary method 2** for detailed procedure).

To investigate the accuracy and precision of the fluorescence-based characterization method, a separate set of AFM stiffness measurements was also performed on the same hydrogel samples (**Figure 3E, 3F, 3G**). Reference marks on the hydrogel-attached coverslips were employed to identify the same location of the gradient region under both measurement modalities. Comparative analysis revealed that the fluorescence-derived stiffness results closely mirrored the AFM measurements in all instances (**Supporting information, Table S1**). Notably, these examples demonstrated that the fluorescence-based characterization technique can effectively predict a wide range of stiffnesses (≈1 to 100 kPa) and gradient slopes (≈10 to 100 kPa mm^-1^), showcasing its potential utility for controlled cell-substrate interaction studies in the context of mechanobiology. Since the fluorescence signal originated from the labeled polymer chains, the images also exhibited spatial fluctuations that reflect the local structural heterogeneities of hydrogels in their native state. This characteristic uniquely differentiates the proposed technique from other established fluorescence-based strategies, and its minimal susceptibility to photobleaching makes it a valuable tool for routine characterization, thus enhancing the reliability of stiffness gradient hydrogels critical for this study.

A key benefit of the fluorescently labeled gradient hydrogels is that they enable rapid validation of the resulting stiffness gradient quality. For example, when fabricating multiple samples for replicate experiments, users can simply inspect the hydrogels under a fluorescence microscope to identify aberrant gradient formation, in which case the gels should be excluded from cell studies. Moreover, conventional AFM indentation workflows often cause damage to the delicate hydrogels during the sample mounting/removal process. Here, the fluorescence-based approach reduces this risk by enabling contactless mechanical stiffness characterization, maintaining sample integrity for subsequent cell culture. More importantly, the fluorescence signal provides an *in situ* means to spatially correlate the local hydrogel properties with the observed cellular responses. Overall, the combination of thermophoretic fabrication and fluorescence labeling strategies overcomes key challenges in implementing alternate stiffness gradient hydrogel platforms, making it a potential solution for scaling gradient hydrogel technology for mechanobiology and tissue engineering applications, where understanding the influence of spatial mechanical cues on cell/tissue behavior is essential.

### 2.4. Fibroblast Migration Toward an Optimal Stiffness Range

Having established the workflow to fabricate FITC-labeled hydrogels with biologically relevant stiffness gradients, the materials were applied to investigate cell-substrate interactions. First, 3T3-L1 fibroblast cells were cultured on F-GM hydrogels with two distinct gradient profiles, including a shallow gradient spanning lower stiffness values (5% F-GM; ≈7-18 kPa) and a steeper gradient across higher stiffness range (10% F-GM; ≈44-106 kPa) (**Figure 4**; **Supporting Information, Figure S5**). The goal was to study the fibroblast mechanoresponses to a change in the absolute stiffness or gradient slope. For all samples collected after 2 h, cell density was comparable across the entire gel surface, indicating that the stiffness gradients did not induce biased cell attachment (**Figure 4A, 4B**; top row). After 24 h on the gradient hydrogels, cells exhibited a steady increase in population toward the stiffer regions on the shallow gradient; conversely, on the steeper gradient, higher cell densities were observed toward the softer regions of the substrate, suggesting a biphasic durotaxis behavior (**Figure 4A, 4B**; bottom row). Specifically, fibroblasts preferentially migrated toward a region of intermediate substrate stiffness (≈20-40 kPa), regardless of whether the cells were exposed to softer or stiffer environments on the same substrate, consistent with our previous study.^[20]^ Interestingly, the migration directionality did not appear to be affected by the slope of the gradient ≈14 kPa mm^-1^ vs. 87 kPa mm^-1^), strongly suggesting that 3T3-L1 fibroblasts in 2D cultures are more mechanosensitive to the absolute substrate stiffness than the mechanical gradient slope.

**Figure 4:**
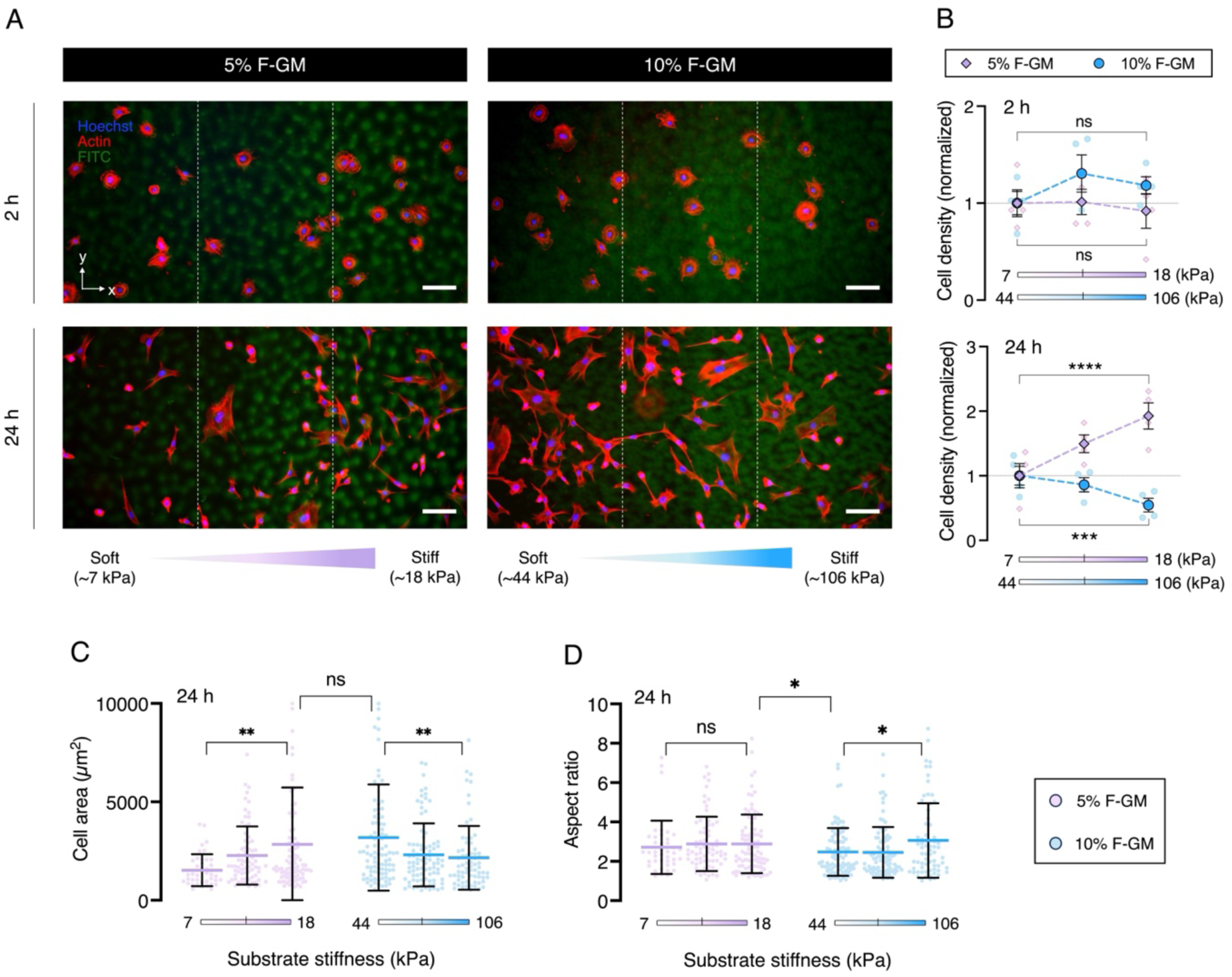
Fibroblast migration on stiffness gradient F-GM is dependent on absolute substrate stiffness. (A) Representative fluorescent images of fibroblasts cultured on shallow (5 wt%) and steep (10 wt%) linear stiffness gradient F-GM hydrogels, observed at 2 h (top) and 24 h (bottom) after seeding. Scale bars: 100 μm. (B) Quantification of cell density along the stiffness gradients at 2 h (top) and 24 h (bottom). Data at each analyzed stiffness region were normalized to the mean cell density at the softest region of the corresponding gel. Data shown as mean ± SEM of n = 4 independent experiments. (C) Quantification of cell area and (D) aspect ratio of fibroblasts along the stiffness gradients at 24 h. Data shown as mean ± SD of n = 50 (5% F-GM, soft), 75 (5% F-GM, intermediate), 101 (5% F-GM, stiff), 94 (10% F-GM, soft), 87 (10% F-GM, intermediate), 71 (10% F-GM, stiff) segmented cells from four independent samples for each gradient system. * (p < 0.05), ** (p < 0.01), *** (p < 0.005), ns (not significant).

To further probe for possible confounding effects of stiffness-dependent cell proliferation, parallel set of experiments were conducted under standard or serum-starved (i.e., suppressed cell division ^[3,45]^) culture conditions (**Supporting Information, Figure S6**). The data revealed a consistent graded cell distribution pattern despite differences in proliferation level between both conditions, confirming that the observations were a direct result of cell migration along stiffness gradients.

The presence of an optimal stiffness for cell migration has recently been reported in various cell types, following the molecular-clutch model which describes directed migration toward an intermediate stiffness region where cell-generated tractions are maximal.^[4,46]^ Since the actin cytoskeleton is central to traction force generation and governs cell morphology,^[47]^ the samples were additionally stained for filamentous actin (F-actin). Quantitative analysis of cell area (**Figure 4C**) revealed a similarly biphasic response with peak spreading around the intermediate stiffness regions, while the aspect ratio (**Figure 4D**) remained mostly consistent across the investigated stiffness range. Therefore, it is postulated that the generation of traction forces by 3T3-L1 fibroblasts directly correlates with cell size to drive stiffness-directed migration. Interestingly, the biphasic stiffness responses observed on these gradient GelMA hydrogels differed from previous reports demonstrating that 3T3 fibroblasts consistently migrate toward stiffer environments, which have traditionally been performed using bioinert polyacrylamide gels functionalized with fibronectin or collagen.^[12,48]^ Together, these findings add to the growing body of evidence that the effect of substrate stiffness on cell behavior is dependent on the composition of matrix to which cells are attached.^[49,50]^

### 2.5. Fibroblast Response to Stiffness Gradients Depends on Matrix Composition

The altered stiffness mechanosensation of cells on different culture substrates has been proposed to be attributed to the distinct material surface properties,^[51]^ such as viscoelasticity,^[52,53]^ wettability,^[54,55]^ and roughness.^[56]^ An active area of research in this respect aims to parse out whether these other factors modulate cell interpretation of substrate stiffness or can independently direct cell behavior. However, systematic comparisons across different biomaterials have proven to be difficult due to constraints in current fabrication strategies, especially in the context of microscale stiffness gradients. To address this challenge, here we further showcase the material flexibility of the thermophoretic fabrication method to investigate the behavior of 3T3-L1 fibroblasts in response to 2D stiffness gradients in F-GG and F-GM systems (**Figure 5**; **Supporting Information, Figure S7**).

**Figure 5:**
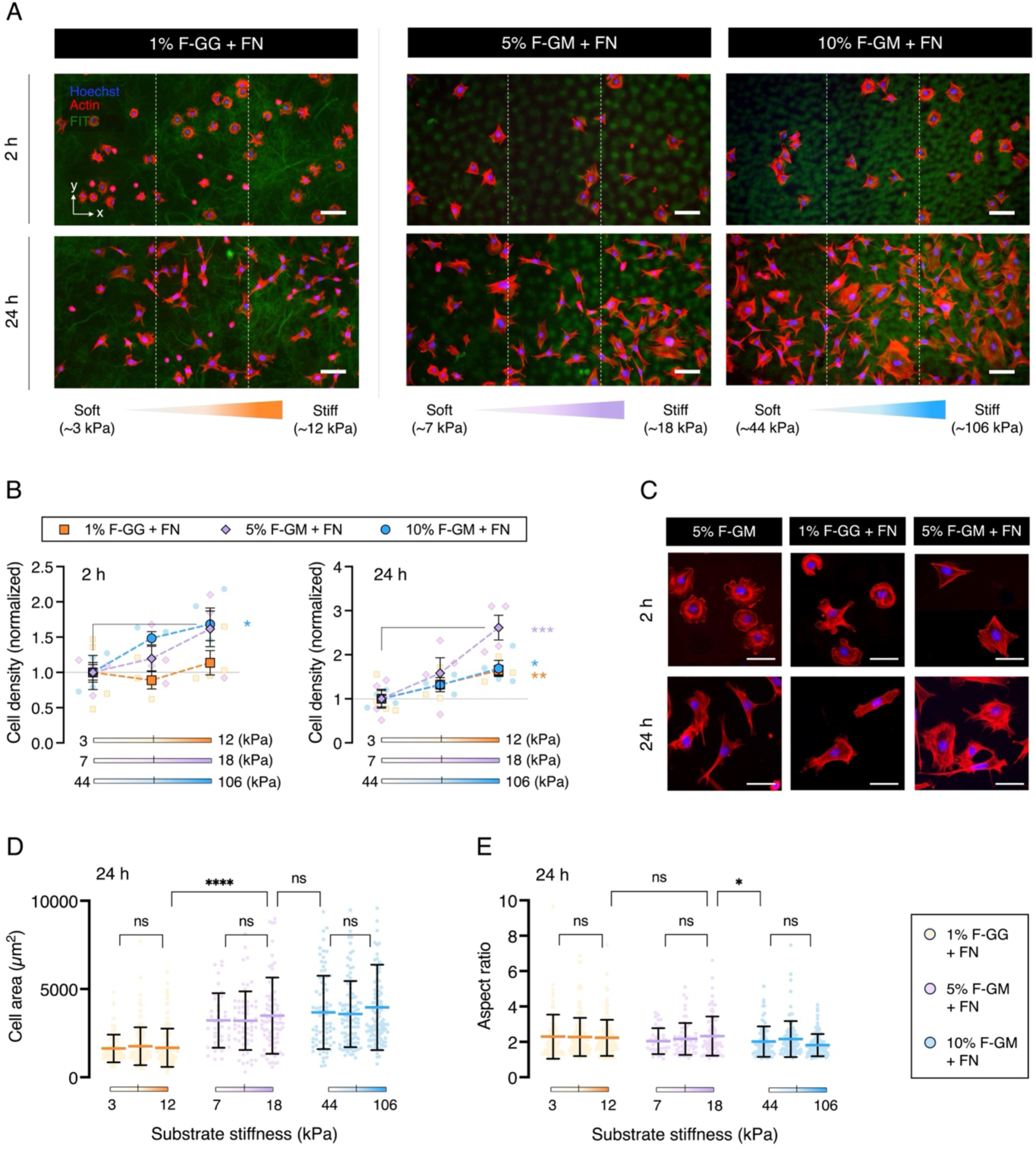
Fibroblast stiffness responses is dependent on the hydrogel matrix composition. (A) Representative fluorescent images of fibroblasts cultured on varying linear stiffness gradient F-GG and F-GM hydrogels with fibronectin (FN) coating, observed at 2 h (top) and 24 h (bottom) after seeding. Scale bars: 100 μm. (B) Quantification of cell density along the stiffness gradients at 2 h (left) and 24 h (right). Data shown as mean ± SEM of n = 4 independent experiments. Data at each analyzed stiffness region were normalized to the mean cell density at the softest region of the corresponding gel. (C) Representative zoom-in fluorescent images of cell morphology on varying linear stiffness gradient F-GG and F-GM hydrogels, observed at 2 h (top) and 24 h (bottom). For consistency, cells were selected from the ‘stiff’ zone on each gradient system. Scale bars: 50 μm. (D) Quantification of cell area and (E) aspect ratio of fibroblasts along the stiffness gradients at 24 h. Data shown as mean ± standard deviation of n = 92 (1% F-GG+FN, soft), 109 (1% F-GG+FN, intermediate), 120 (1% F-GG+FN, stiff), 38 (5% F-GM+FN, soft), 61 (5% F-GM+FN, intermediate), 88 (5% F-GM+FN, stiff), 81 (10% F-GM+FN, soft), 99 (10% F-GM+FN, intermediate), 110 (10% F-GM+FN, stiff) segmented cells from four independent samples for each gradient system. * (p < 0.05), ** (p < 0.01), *** (p < 0.005), ns (not significant).

3T3-L1 fibroblasts were cultured on linear stiffness gradient F-GG and F-GM hydrogels with a similar range (≈5-15 kPa) and slope (≈10 kPa mm^-1^). Unlike GelMA, Gellan gum chains do not contain cell-adhesive domains and thus require surface functionalization with ECM protein to promote cell attachment.^[57]^ For consistency between experiments, both F-GG and F-GM gel samples were conjugated with fibronectin. Furthermore, to assess the impact of fibronectin on cell behavior along F-GM stiffness gradients, the previous experiment using the steep gradient (≈44-106 kPa) was repeated with fibronectin coated using the same procedure. In all cases, immunofluorescent staining to visualize the distribution of fibronectin highlighted spatially uniform functionalization across the stiffness gradients (**Supporting Information, Figure S8**). Thus, this finding provides a confirmation that any difference in the observed cellular responses was not a consequence of differential levels in fibronectin density.

Cells on F-GG gels exhibited no significant difference in cell attachment across the stiffness gradient, while those cultured on F-GM gels showed progressively increasing attachment with substrate stiffness, albeit non-significantly for the shallow gradient (**Figure 5A, 5b**; 2 h). Observations after 24 h further revealed a biased distribution in cell density toward stiffer substrate regions on all gradient hydrogel systems (**Figure 5A, 5b**; 24 h). However, the extent of accumulation was noticeably greater on the shallow gradient F-GM hydrogel than the F-GG hydrogel, supporting the view that the choice of underlying substrate contributes to cell mechanosensing of substrate stiffness. Perhaps even more interesting was that cells on fibronectin-coated F-GM hydrogels did not exhibit a biphasic stiffness-dependent response, but instead displayed a continuously increasing cell population trend over the full ≈7-106 kPa stiffness range (**Supporting Information, Figure S7**). Additionally, cell morphology was found to be influenced by both the ECM protein and base substrate type. Using the fibronectin-coated platform, it was clear that cells showed overall significantly smaller cell size on the F-GG gels compared to the F-GM gels (**Figure 5D**). However, there appeared to be no discernible differences in cell spreading (**Figure 5D**) and aspect ratio (**Figure 5E**) across the stiffness range investigated for each hydrogel type.

Furthermore, single-cell morphology analysis showed that during the initial attachment stage, 3T3-L1 fibroblasts on F-GG and uncoated F-GM substrates displayed an orthoradial pattern of actin filaments in the cell cortex, which is characteristic of immature adhesion (**Figure 5C**; 2 h). In contrast, cells on fibronectin-coated F-GM substrates with the same underlying stiffness profile appeared more well-spread with directionally aligned stress fibers, resembling behavior on ‘stiff-like’ substrates. After 24 h of culture, cells on uncoated F-GM maintained a spindle-like morphology, while fibronectin-coated F-GM promoted rounder cells with enhanced stress fiber formation, whereas cells on F-GG substrates adopted a distinct cobblestone shape (**Figure 5C**; 24 h). These morphological adaptations indicate the importance of cytoskeletal remodeling in governing stiffness-mediated cell responses, which has been well documented to be dependent on the ECM coating mix, and further demonstrated here to have some contributions from the substrate type.^[58,59]^

## 3. Conclusion and Outlook

This study describes a facile solution to the practical challenges associated with the development and deployment of microengineered stiffness gradient hydrogels, particularly for mechanobiology research. The strategy is based on thermophoretic control of polymer distribution in hydrogels created using FITC-labeled polymer chains, offering a material-intrinsic fluorescence readout to probe the local substrate mechanics. Here, photopolymerized GelMA and thermosensitive Gellan gum were used as model systems to demonstrate key advantages in terms of accelerated process optimization, simple gradient verification, and a straightforward means to correlate the local stiffness with cellular responses. The utilization of standard fluorescence microscopy instead of specialized equipment such as AFM underscores the rapidity of material characterization and accessibility of this experimental workflow, which are critical to advance the field of stiffness gradient hydrogels toward more mainstream adoption. Additionally, the fluorescence-based stiffness mapping method demonstrated excellent measurement precision under various stiffness regimes (1-100 kPa) and flexibility with different hydrogel network, validating its reliability for robust biological experiments.

Furthermore, this study emphasizes the importance of various material physiochemical properties in the design of stiffness gradients to elicit specific biological responses. A particularly interesting finding was the synergistic effects of hydrogel type and ECM protein coating on 3T3-L1 fibroblast cell behavior in response to stiffness gradients. Under a similar stiffness range and gradient slope, cells exhibited distinct stiffness-mediated morphological and migration responses when cultured on Gellan gum, GelMA, and fibronectin-coated GelMA platforms. These observations were accompanied by changes in cytoskeletal organization patterns, indicating that the matrix composition might alter cells’ interpretation of substrate stiffness. Future work could leverage similar gradient platforms to probe the exact mechanistic pathway by which different combinations of underlying substrate and ECM protein type could modulate cell mechanoresponses, as well as to assess the generalizability of this interplay across different cell types. To that end, the thermophoresis-based microfabrication technique could be implemented with a wide range of material formulations and programmable gradient patterns, which opens exciting opportunities to gain unique insights into cell-materials interactions that may not be apparent through traditional polyacrylamide or PDMS platforms.

In summary, the method developed here simplifies the experimental complexity and improves the reliability of microengineered stiffness gradient hydrogels for cell culture applications. While the concept of using fluorescence signal as an indicator of hydrogel stiffness is not entirely new, the strategy of directly labelling the hydrogel network is more robust against photobleaching effects than classical approaches based on fluorescein dispersion in the aqueous phase. Although the demonstrations here were focused on FITC for fluorescence labeling due to its low cost and wide availability, the potential for utility with other fluorophore options remains promising. With further development paired with innovations in imaging strategies, this method could likely offer added advantages for researchers by enabling simultaneous microstructure analysis and spatiotemporal tracking for ECM-related studies. For example, real-time visualization of the dynamic cell-substrate remodeling process leading to alterations in the substrate stiffness could be of great interest. Beyond fundamental cell biology research, such comprehensive understanding of the material structure-function relationships can be harnessed toward realizing the goal of designer biomaterials for tissue engineering and regenerative applications.

## 4. Experimental Section

*Preparation of Fluorescently Labeled Polymers*: FITC-GelMA (F-GM) was prepared following a procedure adapted from a previous report.^[31]^ First, GelMA was synthesized by direct reaction of gelatin (Sigma-Aldrich) and methacrylic anhydride (Sigma-Aldrich),^[20]^ resulting in ∼85% modification as confirmed using ^1^H-NMR. 0.4 g of GelMA was dissolved in 10 mL Phosphate buffered saline (PBS) under magnetic stirring at 50 °C. Then, 0.4 mg of FITC (Sigma-Aldrich) dissolved in 400 µL dimethylsulfoxide (DMSO, AJAX FineCHEM) was added to the GelMA solution. The reaction mixture was allowed to mix for 2 h at 50 °C, protected from light. The product was dialyzed against Milli-Q water at 40 °C for 6 days, lyophilized, and stored at –80 °C until use.

FITC-Gellan gum (F-GG) was prepared by adapting previously published protocols on fluorescent labeling of polysaccharides.^[36,37,60]^ Briefly, 1 g of Gellan gum (Sigma-Aldrich) was dissolved in 100 mL DMSO by stirring at 90 °C for 30 min. Once fully dissolved, 10 mg of FITC was added to the solution and allowed to mix for 10 min, after which 80 µL dibutyltin dilaurate (DBTDL, Sigma-Aldrich) was added as catalyst. The reaction was carried out for a further 4 h under continuous stirring and heating at 90 °C, protected from light. The product was washed by three cycles of precipitation in 5× reaction volume of acetone, followed by centrifugation (Eppendorf 5810; 11000 rpm for 5 min) to remove impurities in the supernatant. Finally, the F-GG precipitate was redissolved in Milli-Q water, dialyzed at 40 °C for 6 days, lyophilized, and stored at –80 °C until use.

*Microfluidic Device Fabrication:* Microfluidic devices with configured channels for cooling (water flow) and heating (embedded Joule heaters) were used for imposing local temperature gradients, as described previously.^[19,20]^ All microdevices were fabricated using PDMS (Sylgard 184, Dow Corning) following standard soft lithography techniques. For thermophoresis experiments, the assembled device had a main sample channel (600 µm (width) × 100 µm (depth)) flanked by two co-running side channels (2000 µm (width) × 100 µm (depth)). For gradient hydrogel fabrication, the microdevice was prepared with a pair of heating and cooling channels with cross-sectional dimensions of 1000 µm (width) × 100 µm (depth). An external sample microchamber with 120 µm thickness was made using cut-outs of stacked polymide Kapton tape (Multicomp Pro) and attached to the coverslip-side of the microdevice, forming a complete platform assembly. In both platform designs, the separation between each microchannel was 100 µm.

*Thermophoresis Studies*: Polymer solutions were prepared by mixing FITC-labeled polymer with unlabeled polymer in Milli-Q water in a mass ratio of 1:9. The final GelMA and Gellan gum concentrations were 10 wt% and 1 wt%, respectively. Warm polymer solution was injected into the sample microchannel and left to stabilize for 10 min, ensuring that no flow was present at the beginning of each experiment. Time-lapse images were acquired every 30 s for 40 min. Details of concentration gradient evaluation are provided in **Supplementary method 1**.

*Fabrication of Gradient Hydrogels:* For the F-GM gradient gels, precursor solutions were prepared in PBS at a final concentration of 5 wt% or 10 wt% F-GM and 0.1 wt% Irgacure 2959 (Sigma-Aldrich) as photoinitiator. For the F-GG gradient gels, precursor solutions were prepared in Milli-Q water at a final concentration of 1 wt% F-GG and 5 mM CaCl_2_ (Sigma-Aldrich) as ionic crosslinkers, maintained at 85 °C with gentle agitation to prevent premature gelation.

Linear stiffness gradient hydrogels were fabricated using the procedure detailed in a previous work.^[20]^ Briefly, the microfluidic platform was stabilized at the predefined temperature profile. Then, precursor solution was added to fill the microchamber and immediately covered with a methacrylated coverslip (for F-GM) or poly(ethyleneimine)-coated coverslip (for F-GG) for hydrogel attachment. The applied temperature gradient was maintained for 35 min and 70 min for F-GM and F-GG, respectively. Throughout this process, the device was protected from light to minimize photobleaching. After the indicated duration, F-GM polymerization was achieved by UV exposure (OmniCure S2000 Elite spot curing system, 365 nm, 0.85 W cm^-2^) for 90 s, whereas F-GG crosslinking was achieved by rapid cooling to 15 °C and maintained for 10 min to ensure complete gelation. Finally, the gradient hydrogel was gently peeled off the microchamber, washed twice with PBS, and stored immersed in PBS at 4 °C until use for material characterization or cell study.

*AFM*: Young’s moduli across the stiffness gradient hydrogels were measured using an Asylum MFP-3D AFM. Samples were indented immersed in PBS using a 10 µm diameter borosilicate glass colloidal probe (CP-qp-CONT-BSG, NanoAndMore), with 15 nN trigger force, 2 µm s^-1^ approach velocity, and 10 µm s^-1^ retraction velocity. The cantilever spring constant was calibrated using the thermal noise method at the start of each experiment. The local stiffness at each point was calculated by taking the average of a nanomechanical map generated in a 4 × 4 grid over an area of 20 µm^2^ (16 indentations in total). For gradient samples, a rule with 50 µm intervals printed on high-resolution film photomask (Micro Lithography Services Ltd, UK) was aligned onto the bottom of the sample coverslip to enable precise determination of the x-distance along the stiffness gradient for indentation. The force-displacement curves were analyzed directly in the Asylum MFP-3D software using the Hertz model.

*SEM:* Hydrogel samples were flash frozen using liquid nitrogen and freeze-dried for 1 day. After that, all samples were coated with 10 nm platinum using a Leica ACE600 sputter coater and imaged using a JEOL JSM-IT500HR scanning electron microscope at 5-10 kV. For uniform stiffness samples, images were taken at multiple random locations across the hydrogel surface. For gradient samples, images were taken at regular intervals along the known direction of the stiffness gradient, using the recorded x and y coordinates to locate the soft side (‘soft’), center region (‘intermediate’), and stiff side (‘stiff’) on a sample. The images were then processed and analyzed using FIJI software.

*Confocal Imaging*: Fluorescent images were taken using a Nikon Ti2 confocal microscope at 60× magnification. Hydrogels were imaged directly on the coverslips to which they were fabricated, with a drop of PBS added to prevent sample dehydration during experiments. Details on calibration curve generation and stiffness gradient analysis are found in **Supplementary method 2**.

*Fibronectin coating:* Hydrogels were coated with fibronectin *via* carbodiimide reactions.^[56]^ Samples were washed (2×5 min) with 50 mM MES buffer (2-(N-morpholino)ethanesulfonic acid, pH = 5.5) to prepare the surface for activation. The buffer was aspirated, then the hydrogels were incubated with 100 mM EDC (1-ethyl-3-(-3-dimethylaminopropyl) carbodiimide, Thermo Scientific) and 200 mM NHS (N-hydroxysuccinimide, Thermo Scientific) in MES buffer at room temperature for 30 min. Then, the gels were rinsed twice with PBS to remove unreacted functionalities, and immediately incubated with 100 µg mL^-1^ human plasma fibronectin (Gibco) in PBS at 37 °C for 2 h. Finally, the fibronectin solution was removed, and the gels were washed (2×5 min) with PBS to remove non-specifically adsorbed fibronectin. The coated hydrogel samples were kept immersed in PBS overnight at 4 °C before UV sterilization and cell seeding.

To characterize fibronectin distribution along the stiffness gradients, samples were blocked for 1 h in 1 wt% bovine serum albumin (BSA) at room temperature, then incubated with a rabbit monoclonal anti-fibronectin antibody (ab26820, Abcam, 1:50 dilution) at room temperature for 1 h, followed by secondary goat anti-rabbit IgG antibody (Alexa Fluor 568, A11036, Invitrogen, 1:500 dilution) at room temperature for 30 min.

*Cell Studies:* Mouse fibroblasts (3T3-L1) were cultured in DMEM containing high glucose and pyruvate (Gibco) supplemented with 10% fetal bovine serum (FBS), 100 U mL^-1^ penicillin, and 100 µg mL^-1^ streptomycin. Cells were maintained in a standard humidified incubator at 37 °C in a 5% CO_2_ atmosphere. To suppress proliferation, cells were starved in starvation medium (DMEM containing high glucose and pyruvate supplemented with 1% FBS, 100 U mL^-1^ penicillin, and 100 µg mL^-1^ streptomycin) for 12-15 h before experiments. For experiments, 3T3-L1 cells (up to passage 20) were resuspended in starvation medium and seeded at 3000 cells cm^-2^ in a total volume of 50 µL per hydrogel sample. An even distribution of cells across the entire hydrogel surface was visually confirmed *via* brightfield microscopy at the start of the experiment. The cells were allowed to settle in the incubator for 45 min before topping up with fresh starvation medium. Cell attachment was measured 2 h after seeding, whereas cell migration was assessed after 24h.

*Cell Imaging and Analysis:* Samples were fixed using 4% paraformaldehyde followed by permeabilization with 0.5% Triton X-100 in PBS. Cells were subsequently stained with phalloidin for actin visualization (Sigma-Aldrich) and Hoechst (Sigma-Aldrich) for nuclei quantification. Stained hydrogel samples were imaged using an Olympus IX73 inverted microscope with a Prime BSI Express sCMOS camera (Teledyne Photometrics), acquired at 10× magnification. The region of stiffness gradient was identified using FITC as the reference channel. For each gradient sample, the image was split into 3 equally sized segments for cell analysis. Quantification of cell density, cell area, and aspect ratio was performed using the “Analyze Particles” feature in FIJI software.

*Statistical Analysis:* Data are shown as mean ± standard deviation (SD) or standard error of the mean (SEM) for a minimum of three independent experiments. Statistical analysis was performed on GraphPad Prism (v10) using one-way ANOVA with Tukey post-hoc test. For biological experiments, the numbers of cells analyzed are indicated in the figure legends.

Statistical significance was denoted as * (p < 0.05), ** (p < 0.01), *** (p < 0.005), or ns (non-significant).

## Acknowledgements

We gratefully thank Prof. Chiara Neto for access to the AFM and feedback on the manuscript. We would also like to thank Dr Carmine Onofrillo for insightful discussions on preparation and characterization of FITC-labeled GelMA. S.W.C. acknowledges support from the University of Sydney Faculty of Engineering Research Scholarship and the Paulette Isabel Jones Career Development Award.

## Author Contributions

S.W.C. designed the research, conducted the experiments, analyzed the data, and wrote the original manuscript. L.L. performed SEM experiments and advised on figures development.

D.K. performed confocal imaging. Y.Z. assisted with cell culture experiments. K.K., M.M.M.B., L.A.J., M.B. provided supervision, and acquired resources and funding. D.V. designed the research, provided supervision, and acquired resources and funding. All authors have reviewed and approved the final version of the manuscript.

