## Supplementary Information for "Fluorescently Labeled Gradient Hydrogels Reveal Matrix-Dependent Cell Responses to Substrate Stiffness"

**Supplementary method 1: Fluorescence analysis of polymer concentration profile**

The ability to directly visualize the polymers allowed for material-specific insight into the thermophoretic gradient generation process. In the presence of a temperature gradient, thermophoresis induces directed spatial redistribution of polymers in solution. The result is a progressive formation of a concentration gradient, which can be measured by assessing the fluorescence intensity across the microchannel over time.

For each experiment, image acquisition was automated using the open source µManager software (<https://micro-manager.org/>). Then, the images were processed using a custom MATLAB (Mathworks, R2021b) workflow to analyze the fluorescence intensity profile.

First, the set of images was normalized to the first frame taken at t = 0 min (i.e., immediately before application of temperature gradient), assuming a uniform temperature throughout the sample channel. This step was crucial to 1) compensate for uneven baseline fluorescence that may be caused by static sample defects, and 2) allowed the fluorescence signal to be quantified as relative intensity shifts corresponding to changes in local concentration. Additionally, to account for potential photobleaching, the average image intensity of each frame was calibrated to match that of the first frame. This image processing step reflects mass conservation in the material system, meaning that the observed changes in fluorescence intensity should only be due to polymer redistribution rather than a loss or gain in total polymer content.

Next, a single intensity line profile was computed for each measured time point by averaging along the channel length (i.e., perpendicular to the temperature gradient field). Finally, a linear interpolation was fitted to the central 65% segment of the intensity profile and the slope was extracted, which is directly proportional to the concentration gradient.

**Supplementary method 2: Fluorescence-based stiffness mapping**

By using fluorescently labeled polymers for hydrogel fabrication, it is possible to predict the Young’s modulus of the stiffness gradient hydrogel based on the fluorescence signal measured using a confocal microscope. The core of the workflow consists of two main steps: 1) calibration curve generation and 2) intensity-stiffness correlation, as described below. All image processing and analysis procedures were performed using a customized MATLAB script.

*Calibration curve generation*

For each polymer system, calibration curves connecting the hydrogel concentration to the fluorescence intensity and atomic force microscopy (AFM)-measured stiffness were first generated. To do this, uniform F-GM and F-GG hydrogels were prepared at a series of concentrations. For each sample, 25 µm $\times$ 25 µm fluorescence images were taken at 5 random locations to calculate an averaged pixel intensity, followed by stiffness measurement by AFM (as described in the main methods section). Separate concentration-intensity and concentration-stiffness plots were generated, then correlation equations were determined using simple linear regression. Note that in this study, the fluorescence intensity values were normalized by the intensity at 10 wt% for F-GM and 1 wt% for F-GG, chosen arbitrarily and defined as the reference concentration. This approach was implemented to account for variability in fluorescence readout between different imaging setups, meaning that the established calibration curve equations can be directly applied to subsequent experiments. It should also be noted that while the calibration curves in this study could be sufficiently described by a linear relationship, a different material system or concentration range might require a more complex fitting model; as such, it is recommended that users generate their own set of calibration curves according to the target design space.

*Stiffness gradient hydrogel measurement*

Stiffness analysis was performed by taking a single confocal plane image over the whole gradient region of the thermophoretically fabricated hydrogels. Each pixel intensity was normalized by a reference intensity value, defined as the absolute intensity at the reference polymer concentration (*I_ref_*). Specifically, *I_ref_* should be calculated for each sample image following the equation: *I_ref_* *= Î_img_ / I_N,ref_*, where *Î_img_* is the average intensity of the whole image, and *I_N,ref_* is the normalized intensity at the initial concentration (extracted from the concentration-intensity curve). This step works on the basis that *Î_img_* reflects the absolute intensity at the starting polymer concentration prior to thermophoretic gradient generation, due to mass conservation, as detailed in the previous section. Notably, this is a unique characteristic of the thermophoresis process, and which provides an internal reference point for calibration of each sample. After computing the normalized pixel intensity values, the distribution of the polymer concentration across the gradient region could be determined using the concentration-intensity correlation equation. Finally, the concentration values were connected to the hydrogel stiffness using the concentration-stiffness correlation equation, yielding a heat map for each stiffness gradient.

**Degree of FITC substitution analysis**

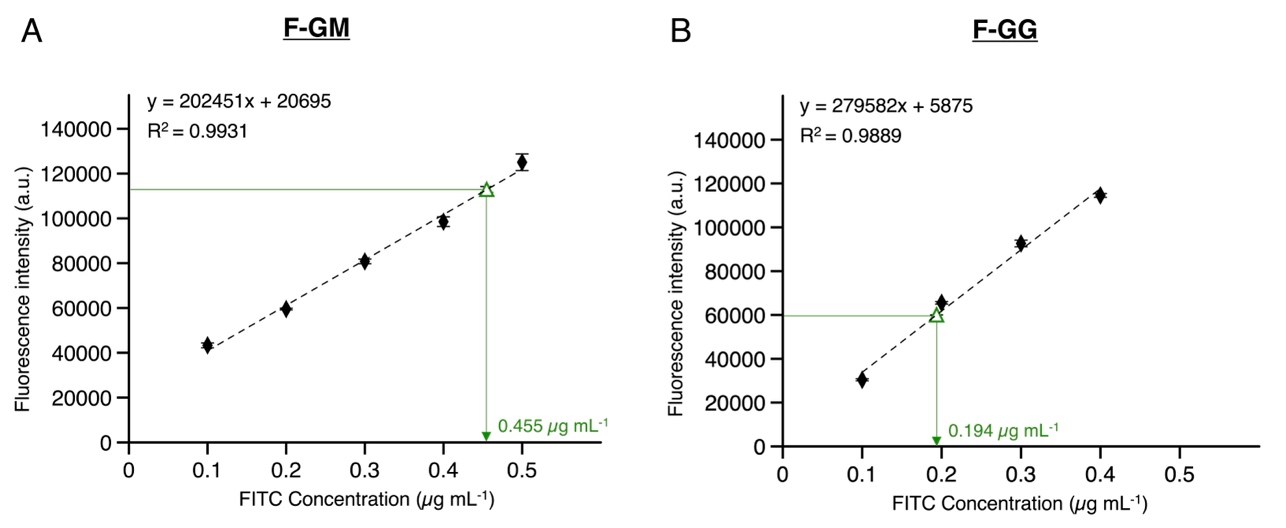

**Figure S1**: The degree of FITC substitution. (A) F-GM and (B) F-GG polymers were characterized using a microplate reader at an excitation wavelength of 483/14 nm and emission wavelength of 530/30 nm (n = 3 independent samples; mean ± SEM). Polymer solutions were prepared in Milli-Q water at a concentration of 50 µg mL^-1^ and compared against a series of FITC standard solution. Water was used as a blank control.

**Effect of FITC labeling on the mechanical and structural properties of hydrogels**

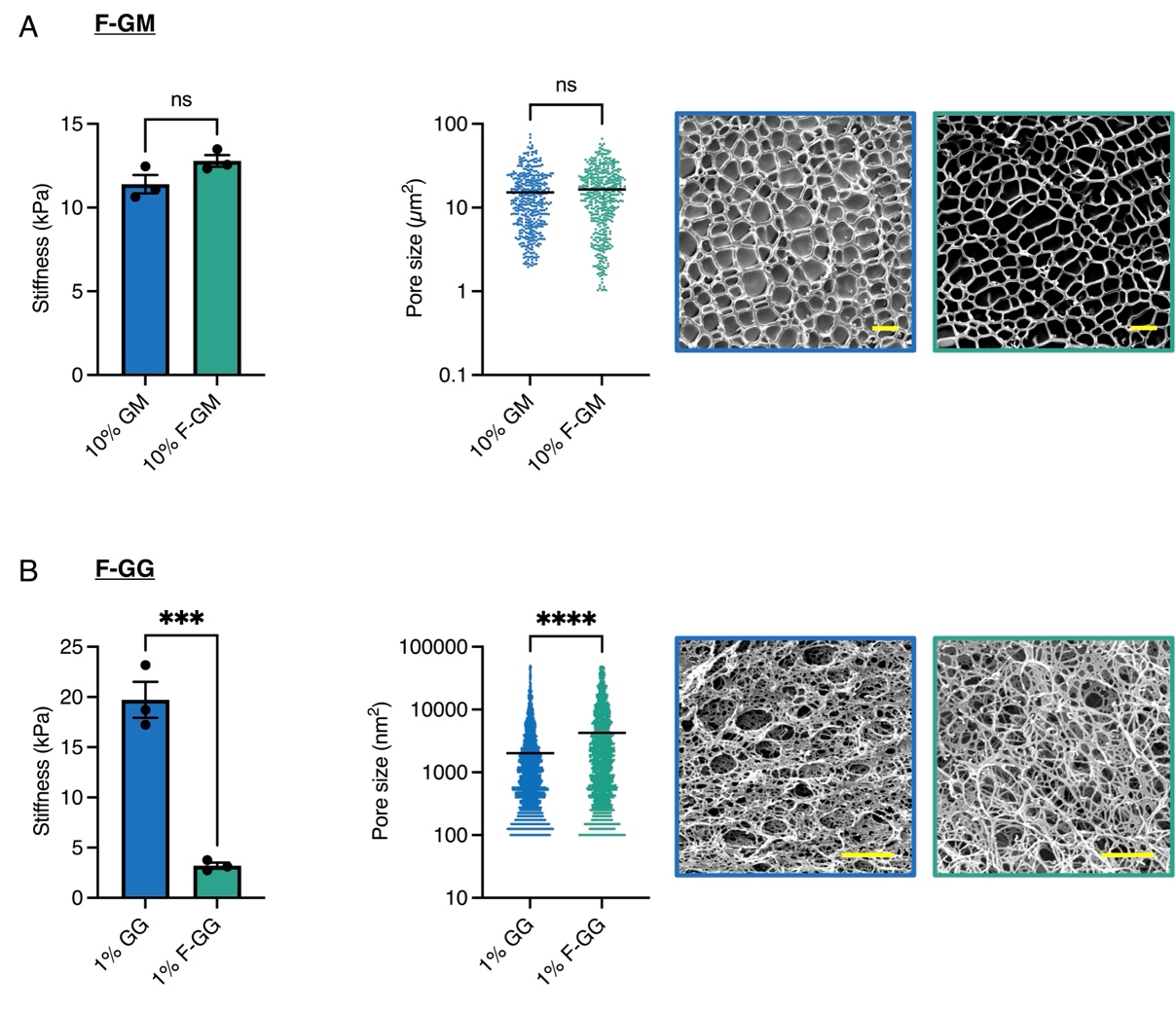

**Figure S2**: AFM stiffness and SEM pore size characterization. (A) 10 wt% GelMA hydrogels made using unmodified polymer (blue) or FITC-labeled polymer (green). Scale bar: 10 µm. (B) 1 wt% Gellan gum hydrogels made using unmodified polymer (blue) or FITC-labeled polymer (green). Scale bar: 1 µm. Stiffness data are shown as mean $\pm$ SEM of n = 3 independent samples. Quantification of pore size is the mean of n = 3 independent samples, where each point depicts an individual pore analyzed. Note that the stiffness of GelMA samples is overall lower than that reported in Figure 1 in the main text due to different UV exposure settings used in these preliminary experiments. * (p < 0.05), ** (p < 0.01), *** (p < 0.005), ns (not significant).

**Thermophoresis characteristic time analysis**

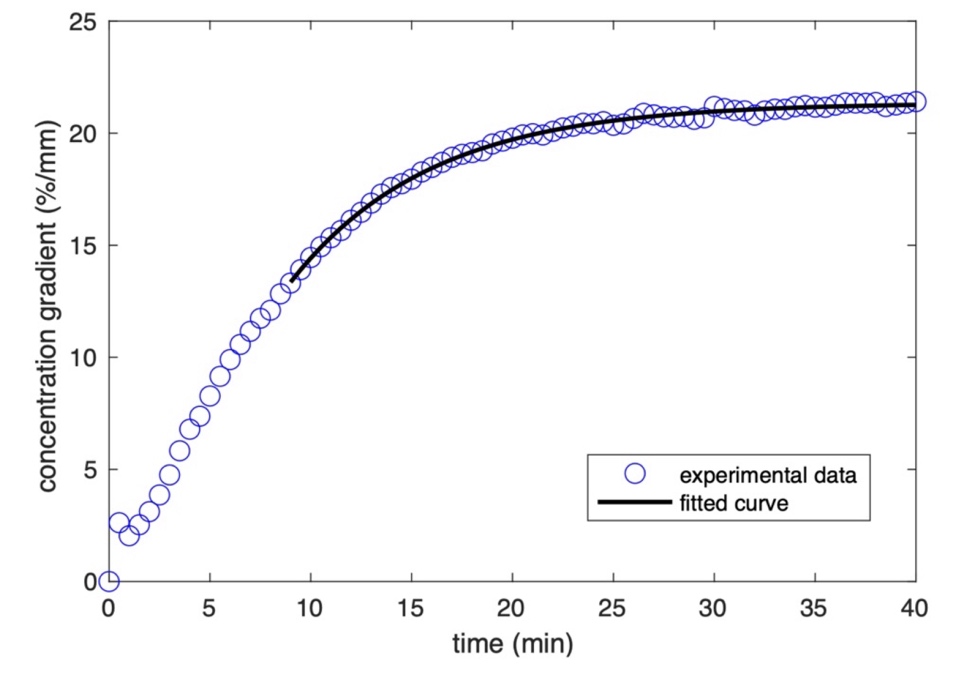

| **Polymer**  **type** | **Temperature gradient**  **[°C mm^-1^]** | $\boldsymbol{\tau}$  **[min]** | **R^2^**  **value** |
| --- | --- | --- | --- |
| GelMA | ≈7 | ≈7 | 0.9975 |
| GelMA | ≈13 | ≈9 | 0.9940 |
| Gellan gum | ≈15 | ≈45 | 0.9825 |
| Gellan gum | ≈18 | ≈16 | 0.9946 |

**Figure S3**: Thermophoresis characteristic time ($\tau$) analysis: (Top) example demonstrating exponential fit [$\nabla c=A-B exp(-t/\tau$)] to the concentration gradient ($\nabla c$) vs. time ($t$) curve, where $A$ and $B$ are constants. Note that the fit started from approximately 1/3 of $\tau$ (estimated from the trajectory of the full plot), to discount potential noise or instability in the initial stages of the process. (Bottom) Summary of $\tau$ evaluated from the curves in Figure 2E in the main text. R^2^ values are reported as a measure of goodness of curve fit.

**Operational conditions for the fabrication of stiffness gradient hydrogels**

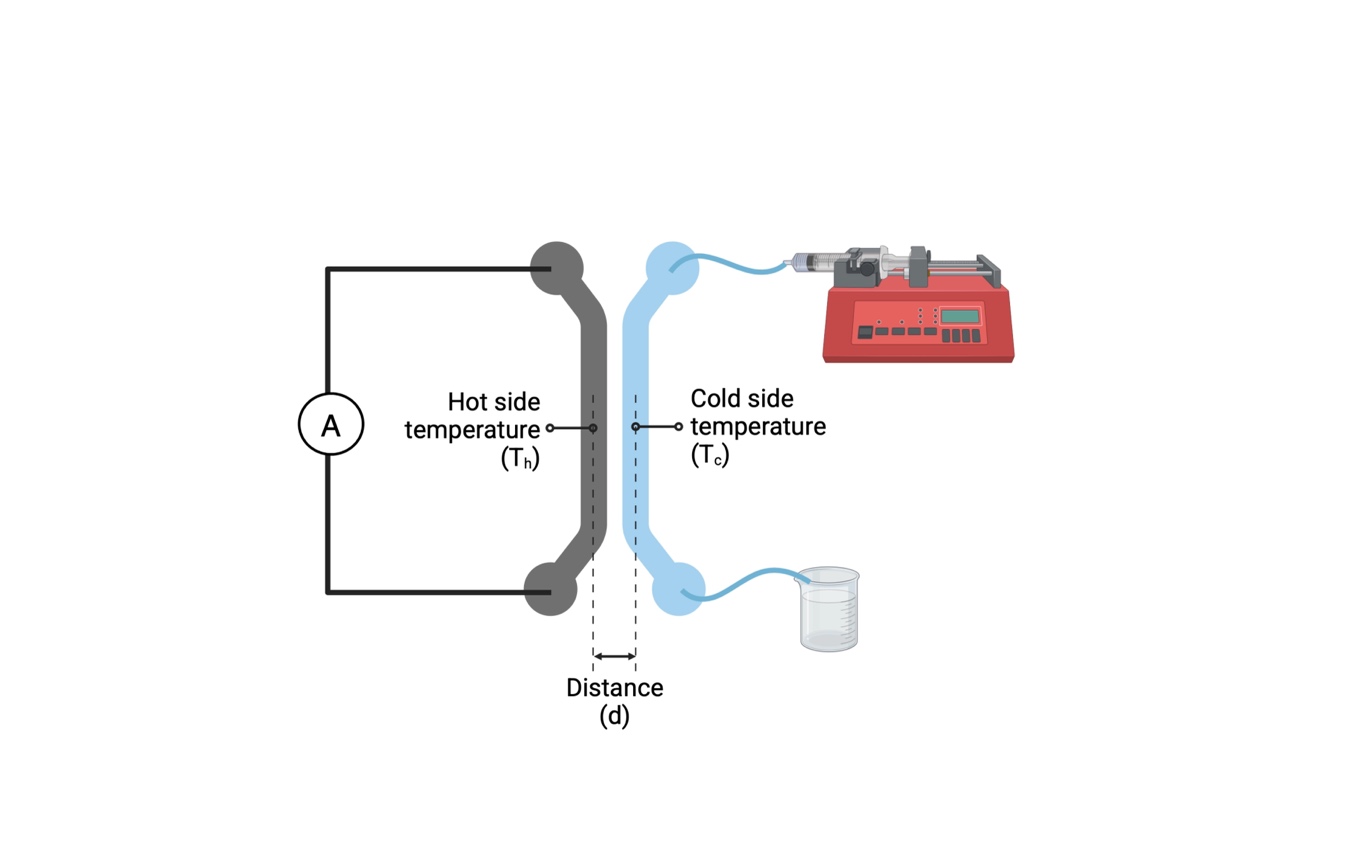

| **Hydrogel type** | **Water**  **flow rate**  **[µL min^-1^]** | **Controlled current**  **[A]** | **Temperature range**  **[°C]** | **Temperature gradient**  **[°C mm^-1^]** | **Process**  **time**  **[min]** |
| --- | --- | --- | --- | --- | --- |
| F-GM | 100 | 1.0 | T_c_ ≈55 / T_h_ ≈65 | 9 | 35 |
| F-GG | 90 | 1.1 | T_c_ ≈55 / T_h_ ≈75 | 15 | 70 |

**Figure S4**: Schematic of the thermal microfluidic system comprising a pair of Joule heating (grey) and water-cooling (blue) channels, used to generate a controlled linear temperature gradient across the fabrication platform. The Joule heater was operated using a power supply unit with controlled DC current, whereas the water flow rate was set using a syringe pump. Different process parameters were used for F-GM and F-GG hydrogels, noting that the applied temperature gradient was defined as [(T_h_-T_c_)/d].

**Fibroblast migration on F-GM hydrogels with linear stiffness gradients**

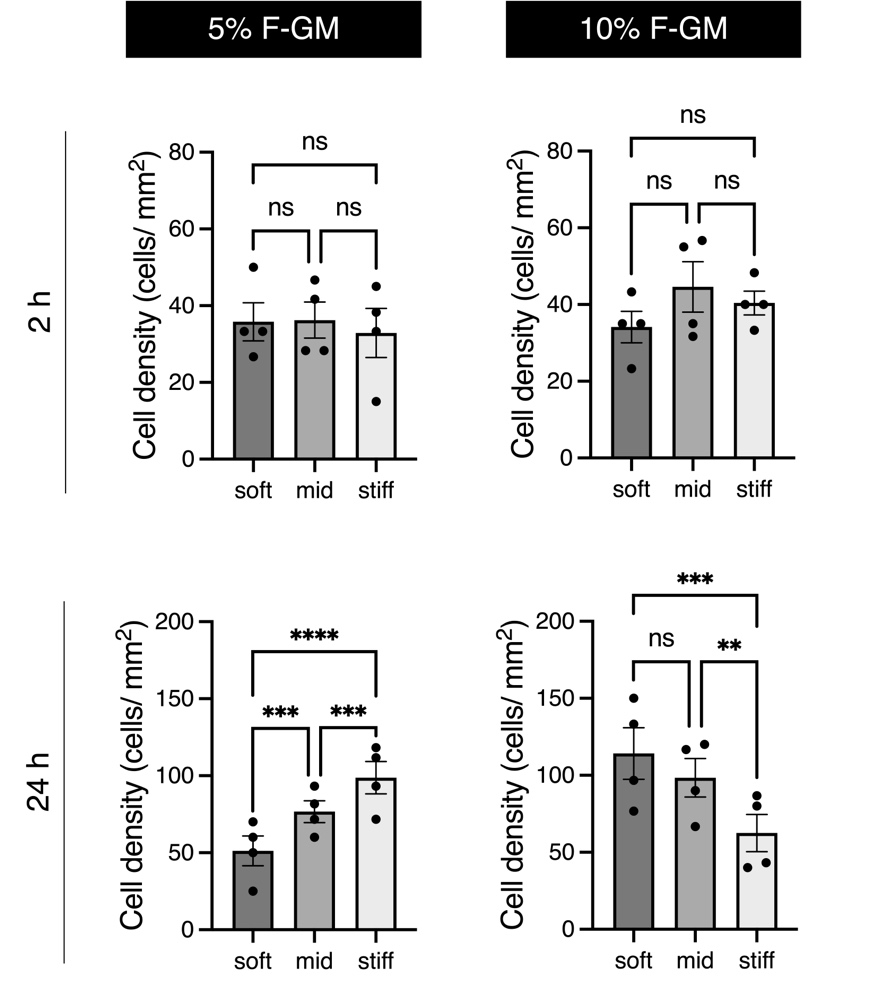

**Figure S5**: Fibroblast cell distribution on shallow (5%) and steep (10%) linear stiffness gradient F-GM hydrogels. Cell density quantified by mean nuclei per analyzed region (‘soft’, ‘mid’, ‘stiff’) after 2 h (top) and 24 h (bottom) culture, corresponding to the normalized data reported in Figure 4B in the main text. Data shown as mean $\pm$ SEM of n = 4 independent experiments. * (p < 0.05), ** (p < 0.01), *** (p < 0.005), ns (not significant).

**Comparison of cell behavior under standard vs. serum-starved culture conditions**

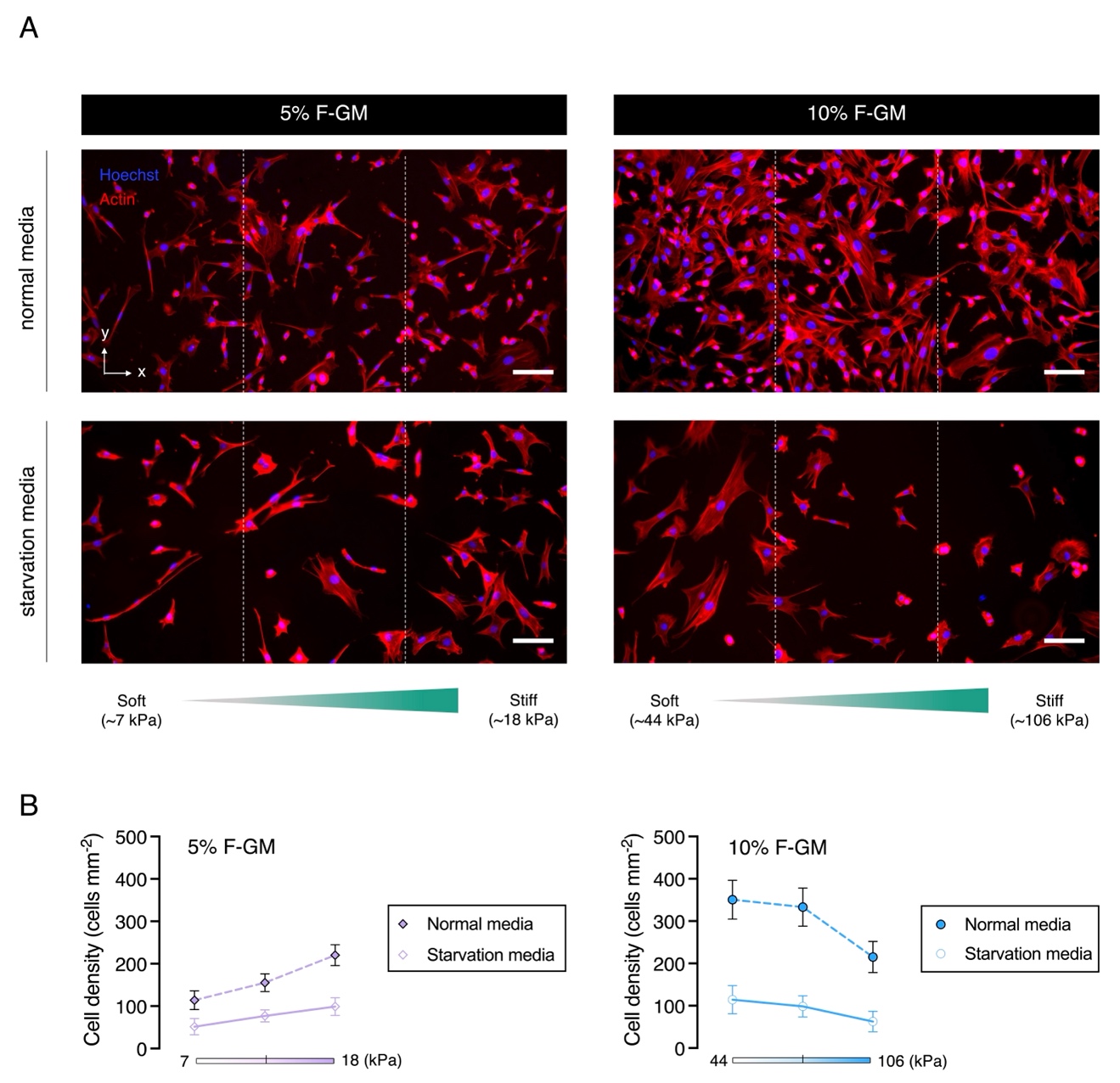

**Figure S6**: Fibroblasts exhibit consistent migration pattern in both culture conditions but lower proliferation activity using serum-starved media. (A) Representative fluorescent images of 3T3-L1 fibroblasts cultured on shallow (5%) and steep (10%) linear stiffness gradient F-GM hydrogels at 24 h, using normal (top) or starvation (bottom) media. Scale bars: 100 µm. (B) Corresponding quantification of cell density along the stiffness gradients. Data shown as mean $\pm$ SEM of n = 4 independent experiments.

**Fibroblast migration on fibronectin coated F-GG and F-GM hydrogels with linear stiffness gradients**

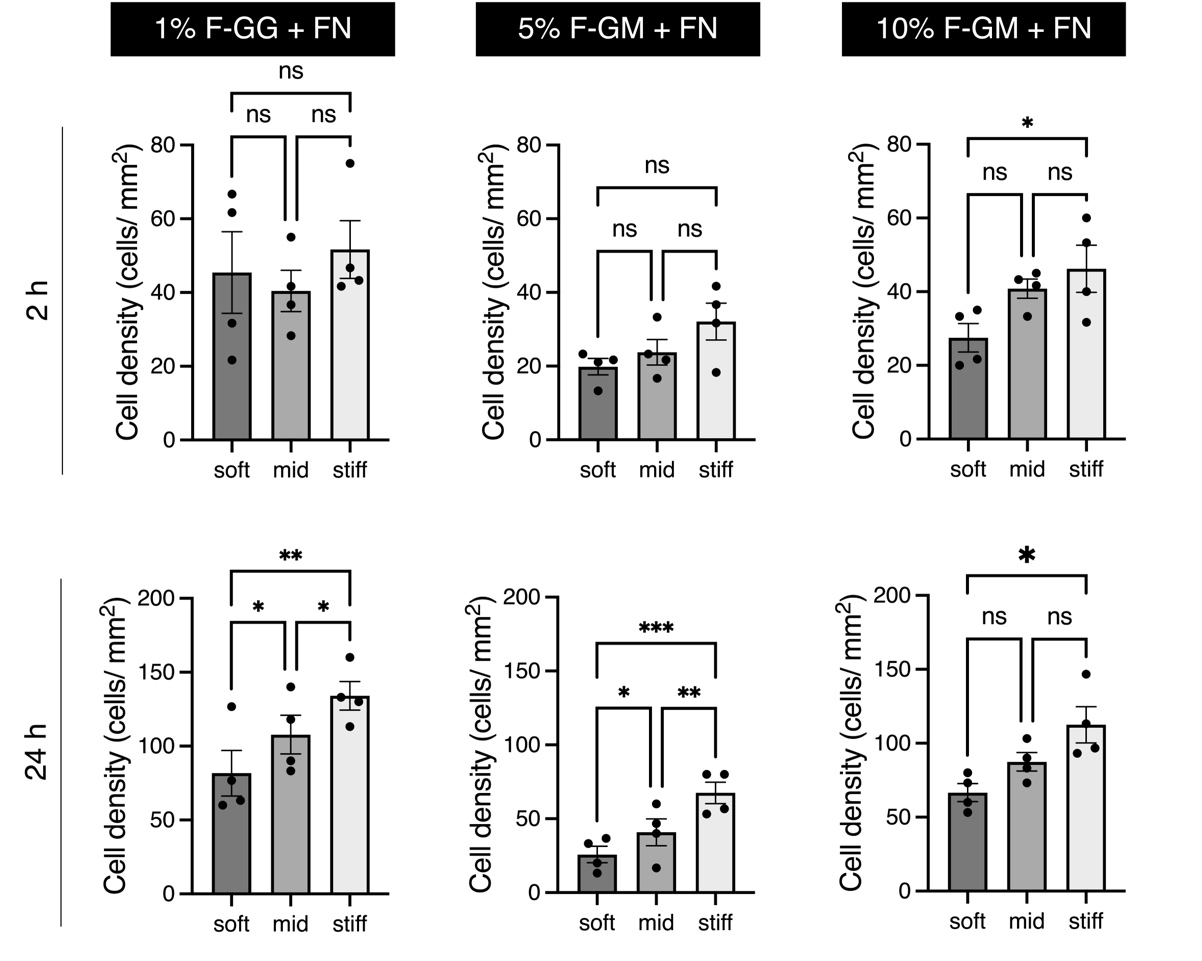

**Figure S7**: Fibroblast cell distribution on varying linear stiffness gradient F-GG and F-GM hydrogels with fibronectin (FN) coating. Cell density quantified by mean nuclei per analyzed region (‘soft’, ‘mid’, ‘stiff’) after 2 h (top) and 24 h (bottom) culture, corresponding to the normalized data reported in Figure 5B in the main text. Data shown as mean $\pm$ SEM of n = 4 independent experiments. * (p < 0.05), ** (p < 0.01), *** (p < 0.005), ns (not significant).

**Fibronectin distribution on thermophoretically fabricated stiffness gradient hydrogels**

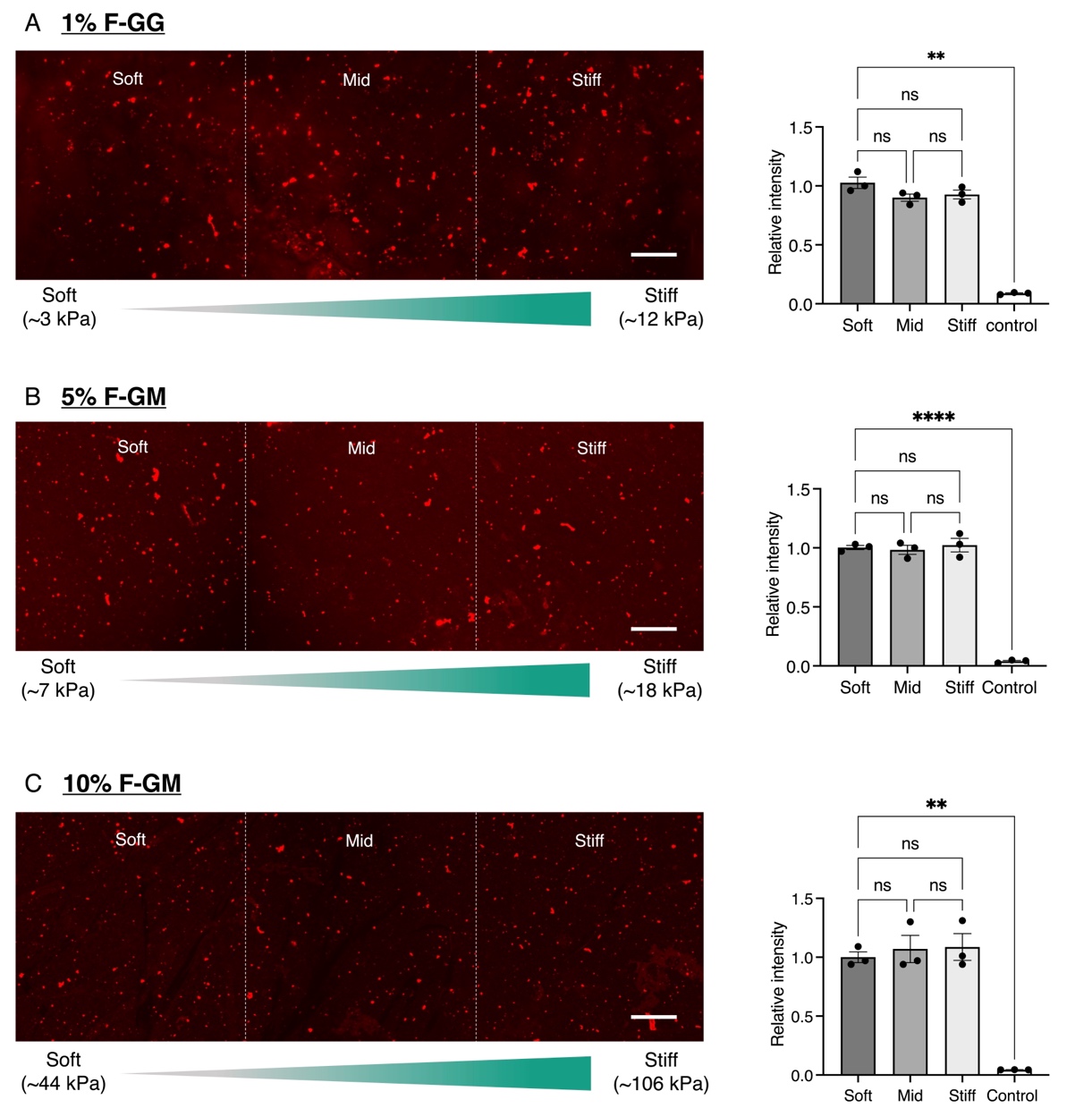

**Figure S8**: Quantification of fibronectin attachment on the hydrogel surface. Representative fluorescent images of linear stiffness gradient hydrogels fabricated using (A) 1% F-GG, (B) 5% F-GM, and (C) 10% F-GM polymer systems (n = 3 independent samples; mean ± SEM). Data were normalized by dividing the sum of fluorescence intensity at each stiffness region (‘soft’, ‘mid’, ‘stiff’) by that at the corresponding ‘soft’ region. Scale bars: 100 µm. Negative control samples were prepared using uniform stiffness hydrogels (1% F-GG, 5% F-GM, or 10% F-GM) without fibronectin coating. * (p < 0.05), ** (p < 0.01), **** (p < 0.005), ns (not significant).

**Comparison of fluorescence-based vs. AFM stiffness measurements**

**Table S1**: Summary table comparing the stiffness gradient properties obtained using the fluorescence-based method, analyzed using the data from Figure 3 in the main text.

| **Gradient hydrogel** | **Stiffness range**  **[kPa]** | | **Gradient slope**  **[kPa mm^-1^]** | |
| --- | --- | --- | --- | --- |
|  | **Fluorescence** | **AFM** | **Fluorescence** | **AFM** |
| 10% F-GM | Low: 44 $\pm$ 7  High: 106 $\pm$ 3 | Low: 42 $\pm$ 5  High: 137 $\pm$ 20 | ≈87 | ≈108 |
| 5% F-GM | Low: 7 $\pm$ 1  High: 18 $\pm$ 1 | Low: 6 $\pm$ 1  High: 15 $\pm$ 2 | ≈14 | ≈10 |
| 1% F-GG | Low: 3 $\pm$ 1  High: 12 $\pm$ 1 | Low: 4 $\pm$ 1  High: 11 $\pm$ 2 | ≈9 | ≈8 |
